# Homologous recombination mutant lethality differs between *h*− and *h+ Schizosaccharomyces pombe* strains due to *mat1* heterochromatin

**DOI:** 10.1101/2025.10.13.682094

**Authors:** Peter Kolesar, Siarhei Paliavoi, Barbora Stefanovie, Jan Josef Palecek

## Abstract

Homologous recombination (HR) is generally considered dispensable in yeast and vertebrates, yet mounting evidence indicates that its essentiality depends on cellular context. Here, we dissect the basis of this context dependency in *Schizosaccharomyces pombe*. In the homothallic *h*^*90*^ strain, regarded as wild type, mating-type switching (MTS) occurs every other cell division and requires HR to repair programmed double-strand breaks (DSBs) at the *mat1* locus. We show that the widely used heterothallic *h*^−*S*^ strain is likewise dependent on HR for viability. HR-deficient *h*^−*S*^ mutants (*rad51Δ, rad52Δ*, or *rad54Δ*), still frequently employed in the literature, survive only when carrying secondary suppressor mutations that abolish *mat1* DSB formation, such as *smt-0, swi1Δ*, or *fml1Δ*. In contrast, HR is dispensable in the *h*^*+N*^ strain, where duplication of the *mat2/3* region into *mat1* introduces the *cenH* and *REIII* elements. These elements nucleate H3K9 methylation and heterochromatin spreading across the imprint site, blocking imprintosome recruitment and thereby preventing both imprinting and DSB formation. Disruption of this heterochromatin, via deletion of *cenH* or key chromatin modifiers, restores DSB formation in *h*^*+N*^ cells and reinstates HR essentiality in the absence of the Clr4 methyltransferase. Collectively, our findings demonstrate that HR is indispensable for *S. pombe* survival due to its critical role in repairing *mat1* DSBs, except under genetic or epigenetic conditions that suppress their formation.

## INTRODUCTION

The fission yeast *Schizosaccharomyces pombe* spends most of its life cycle in the haploid state and exists as one of two mating types, minus (M) or plus (P). Mating occurs only between cells of opposite types. Depending on the stability of mating type inheritance, *S. pombe* strains are classified as heterothallic or homothallic. Heterothallic strains maintain a stable mating type (either M or P) across cell divisions, whereas homothallic strains undergo mating-type switching (MTS), generating populations that contain both P and M cells in similar proportions (reviewed in (1)).

The mating type (*mat*) locus resides on the right arm of chromosome II. In the homothallic reference strain *h*^*90*^, this region contains three related cassettes of ∼1 kb each: *mat1* (harboring either the M or P allele), *mat2-P*, and *mat3-M* (Figure 1A). The P cassette encodes the *Pc* and *Pi* genes, whereas the M cassette encodes the *Mc* and *Mi* genes. All four genes specify mating-type-specific transcription factors, with “c” denoting constitutive and “i” denoting inducible expression. Among these, only *mat1* is transcriptionally active and determines the cell’s mating type. In contrast, *mat2-P* and *mat3-M* remain transcriptionally silent, residing within a heterochromatic domain that is separated from *mat1* by the 17 kb L region. Between *mat2-P* and *mat3-M* lies the 11 kb K region, which contains the 4.3 kb *cenH* repeat array, composed of *dg* and *dh* sequences that nucleate heterochromatin (2). Transcription of these repeats produces noncoding RNAs, which are processed by the RNA interference machinery into small interfering RNAs that recruit the Clr4 methyltransferase to establish H3K9 methylation (H3K9me; (3)). The integrity of the heterochromatic region is reinforced by the *REII* and *REIII* repressor elements (Figure 1A), which recruit histone deacetylases required for subsequent H3K9me. *REII* acts through Clr5, while *REIII* contains binding sites for the Atf1/Pcr1 transcription factors that recruit Clr6 and Clr3 (1,4,5). This heterochromatic domain is flanked by two boundary elements, *IR-L* and *IR-R*, consisting of 2 kb inverted repeats (6).

**Figure 1.**
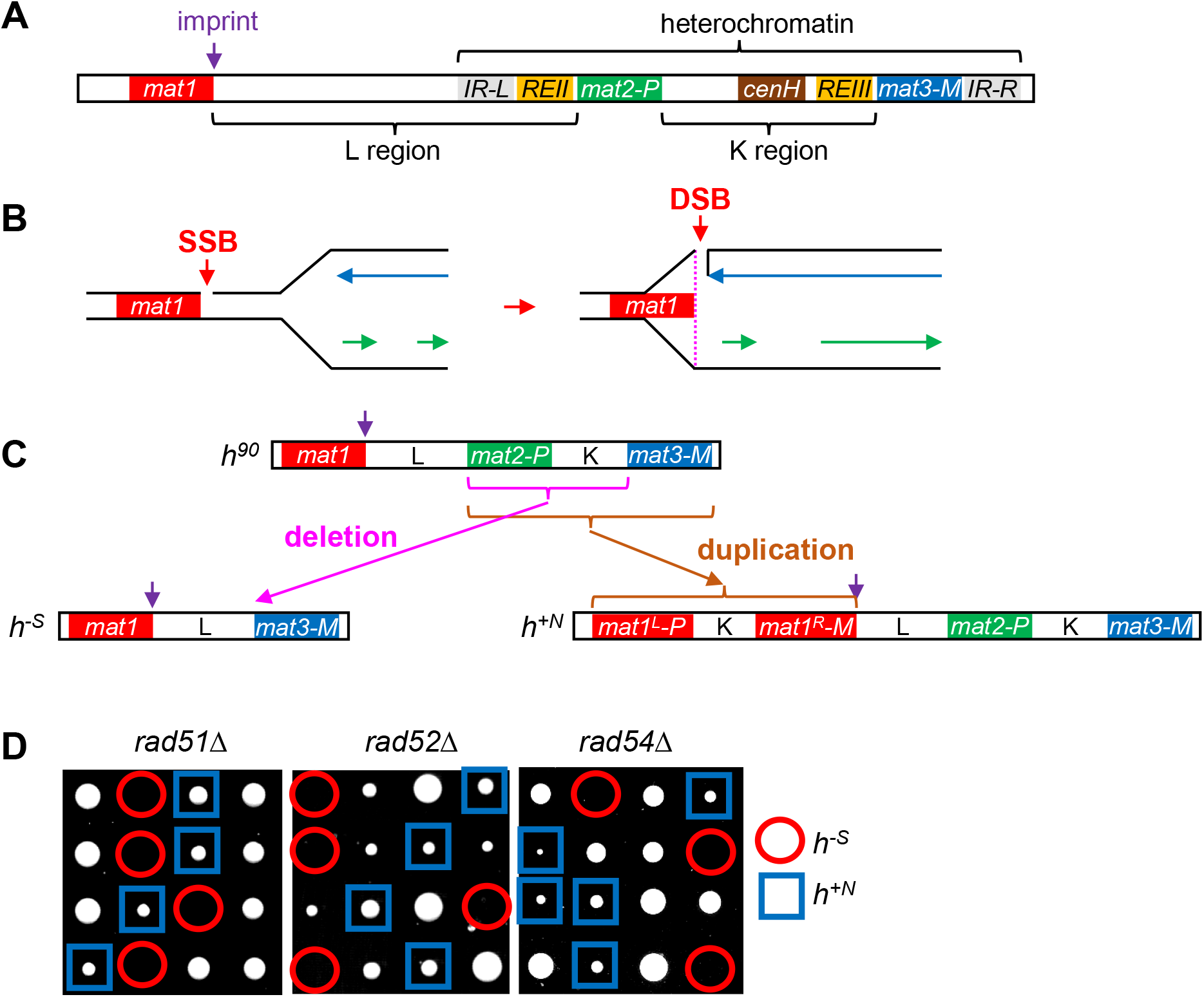
Structure of the *mat* locus and requirement of HR in the *h*^*-S*^ strain. (**A**) Schematic of the *mat* locus on chromosome II showing the major functional elements described in the text. (**B**) During DNA replication, the site-specific imprint at *mat1*, represented as a single-strand break (SSB), is converted into a one-ended double-strand break (DSB) by leading-strand synthesis. (**C**) Organization of the *mat* locus in *h*^*90*^, *h*^*-S*^, and *h*^*+N*^ strains. The *h*^*-S*^ strain arose from *h*^*90*^ through deletion of *mat2-P* and the intervening K region, while *h*^*+N*^ carries an insertion of the entire *mat2*-K-*mat3* sequence into *mat1*. (**D**) HR is essential in the *h*^*-S*^ background. *h*^*+N*^ strains carrying deletions of *rad51, rad52*, or *rad54* were crossed with wild-type *h*^*- S*^, and meiotic progeny were analyzed by tetrad dissection. Mutants in the *h*^*-S*^ background are circled; mutants in *h*^*+N*^ are boxed.

During DNA replication in homothallic strains, MTS is initiated by a site-specific imprint, believed to be either a single-strand break or a ribonucleotide insertion, introduced at the centromere-distal end of *mat1* (Figure 1A and B; (1)). In the subsequent cell cycle, this imprint is converted into a one-ended double-strand break (DSB) by DNA polymerase ε (7). The DSB then initiates gene conversion by invading the silent donor alleles: P cells preferentially use *mat3-M*, and M cells use *mat2-P*, thereby switching their mating type (8).

The homothallic strain *h*^*90*^ is the most common in nature and is considered the wild-type state (9). Heterothallic strains arise spontaneously from *h*^*90*^ via duplications or deletions within the *mat* region (10). The two most widely used laboratory strains are *h*^*-S*^ (M type) and *h*^*+N*^ (P type). The *h*^*-S*^ strain carries a deletion of *mat2-P* and the K region (Figure 1C). While capable of generating the imprint-induced DSB, *h*^*-S*^ lacks an appropriate donor allele for gene conversion, thereby stabilizing the M type. The *h*^*-S*^ (972) genome was sequenced during the *S. pombe* genome project and serves as the reference DNA sequence (11). By contrast, the precise sequence of *h*^*+N*^ remains unpublished, but restriction analyses suggest that *mat1* contains a duplication of silent cassettes (Figure 1C; (10)). *h*^*+N*^ cells retain the P type because the stable left-hand cassette (centromere-proximal; commonly referred to as *mat1:2* in the literature) appears to dictate mating type, whereas the potentially switchable right-hand cassette (*mat3:1*) remains phenotypically silent (10). Additionally, *h*^*+N*^ shows a marked deficiency in *mat1* DSB formation, although the underlying mechanism remains unclear (10,12).

MTS is intimately linked to homologous recombination (HR), the main pathway in fission yeast for repairing DSBs and restarting collapsed replication forks (13). In budding yeast, HR-deficient mutants are viable in heterothallic but not homothallic strains, as DSB formation relies on the HO endonuclease active only in homothallic cells (14). In *S. pombe*, however, HR essentiality has been less clear. According to PomBase, the key database for the fission yeast community, deletion mutants of core HR genes such as *rad51, rad52*, and *rad54* are viable (15). Yet, these data are derived from *h*^*+N*^ strains (16–18), which are inefficient at generating *mat1* DSBs. Already in 1993, Ostermann et al. reported that *rad52* is essential in the homothallic strain *h*^*90*^ but dispensable in the heterothallic *h*^*+N*^ and *h*^*-S*^ backgrounds (19). This was later extended to *rad51Δ* and *rad54Δ*, both of which are lethal in *h*^*90*^ (20). The likely explanation is that HR is indispensable for repairing *mat1* DSBs, which otherwise lead to cell death (1,20). Interestingly, Roseaulin et al. further demonstrated that *rad51Δ* is lethal not only in *h*^*90*^ but also in a heterothallic *mat1-M* strain that lacks donor cassettes (20). This observation suggested that HR is also essential in the commonly used *h*^*-S*^ strain, contrary to the earlier report. Nonetheless, approximately half of recent studies rely on *rad51* or *rad52* deletions in *h*^*-S*^ (Table S1).

In this study, we resolve the longstanding ambiguity surrounding HR essentiality in heterothallic strains. We show that, in contrast to *h*^*+N*^ cells, HR is indispensable for survival in *h*^*-S*^ strains unless secondary mutations alleviate this requirement. Furthermore, we demonstrate that duplication of the *cenH* and *REIII* heterochromatic elements within *mat1* of *h*^*+N*^ cells drives H3K9me across the imprint site. The resulting heterochromatin not only silences local transcription but also blocks imprintosome recruitment, thereby preventing DSB formation and enabling the survival of HR-deficient mutants.

## RESULTS

### HR is indispensable for viability in the *h*^*-S*^ strain

Previous studies established that deletion of the main HR genes *rad51, rad52*, or *rad54* results in viable mutants in the heterothallic *h*^*+N*^ strain, but inviability in the homothallic *h*^*90*^ strain (16–20). Whether these genes are essential in the heterothallic *h*^*-S*^ strain, however, has remained unclear ((19,20); Table S1). To directly address this question, we crossed *rad51*Δ, *rad52*Δ, and *rad54*Δ mutants in the *h*^*+N*^ background with *h*^*-S*^ and analyzed meiotic progeny by tetrad dissection. In each case, *h+* spores carrying the deletions formed smaller but viable colonies, whereas *h-* spores arrested growth shortly after plating and failed to produce visible colonies (Figure 1D). These results demonstrate that, as in *h*^*90*^, HR is indispensable for viability in the *h*^*-S*^ background.

### Viability of HR mutants in *h*^*-S*^ depends on secondary suppressor mutations

If *rad51, rad52*, or *rad54* deletions are lethal in *h*^*-S*^, how can they frequently appear in this background in published studies (Table S1)? The most likely explanation is that these strains carry additional suppressor mutations that bypass the need for HR. Since HR is essential for repairing *mat1* DSBs, preventing DSB formation would restore viability. A common mutation that blocks *mat1* DSB formation is *smt-0* (switch mating type), a 262 bp deletion adjacent to the imprint site of the *mat1* locus (21,22). As expected, *rad51Δ, rad52Δ*, or *rad54Δ* mutants are viable in the *h*^*-S*^ *smt-0* background (Figure 2A). Similarly, deletion of Swi1, required for efficient *mat1* DSB formation (1,23), also suppressed HR mutant lethality (Figure 2B). Combining *swi1Δ* with *rad52Δ* resulted in a synthetic growth defect, manifesting as markedly smaller colonies in both *h*^*+N*^ and *h*^*-S*^ backgrounds. This defect likely arises from the requirement for Rad52-mediated single-strand annealing to repair single-stranded DNA gaps that accumulate in *swi1Δ* cells (24).

**Figure 2.**
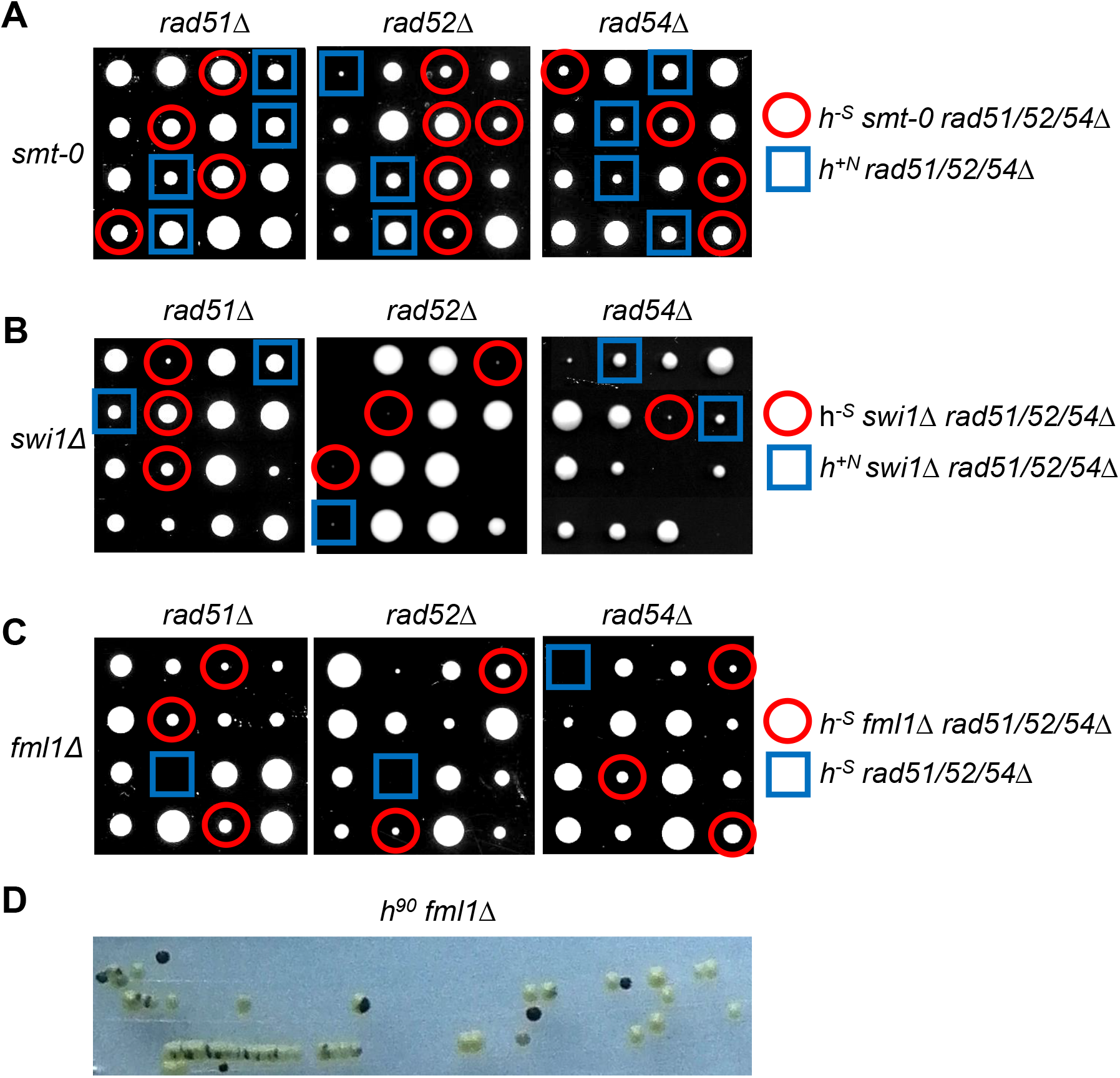
Suppressor mutations that prevent *mat1* DSB formation rescue HR mutant lethality in *h*^*-S*^. (**A-C**) *h*^*+N*^ strains carrying deletions of *rad51, rad52*, or *rad54* were crossed with *h*^*-S*^ strains harboring *smt-0* (A), *swi1Δ* (B), or *fml1Δ* (C) mutations, and meiotic progeny were analyzed by tetrad dissection. (**D**) Fml1 is required for MTS. Iodine staining of *h*^*90*^ *fml1Δ* cells grown on EMM–N plate for 4 days at 28 °C.

To identify suppressors in published strains, we analyzed *h-rad51Δ* strains from the NBRP library (YGRC, Japan). We found that FY18725 was actually Rad51 wild-type, and FY20076 was *h+*. Sequencing FY18537 (*h-rad51::hygr*), used in multiple publications (25–29), revealed additional mutations (File S1), including *cdc6-P313S* (polymerase δ), *rad21-T357N* (cohesin) and *cnd3-S9F* (condensin), likely inhibiting *mat1* DSB formation. Proper regulation of Pol δ’s strand displacement activity is required to prevent complete replacement of the Pol α-synthesized primer, thereby stabilizing the imprint. Concurrently, cohesin and condensin preserve the structural integrity of the *mat* locus, and their importance for MTS has been documented in *S. pombe* and *S. cerevisiae* (30–32).

In another *h-rad51Δ* strain (S7068, (33)), we identified a 330 kb duplication ending just upstream of *fml1*. We hypothesized that this duplication disrupted Fml1 function, and introducing *fml1Δ* into the main HR mutants suppressed their lethality in *h*^*-S*^ (Figure 2C), presumably again by blocking DSB formation. To test whether Fml1 affects MTS, we examined *h*^*90*^ *fml1Δ* strains by iodine staining. Wild-type *h*^*90*^ colonies stain almost black due to high spore content after MTS, whereas MTS mutants stain lighter. *h*^*90*^ *fml1Δ* colonies showed a high proportion of non-stained colonies (Figure 2D), confirming Fml1’s role in MTS. Together, these results demonstrate that *rad51, rad52*, and *rad54* deletions are viable in *h*^*-S*^ only when a secondary mutation suppresses *mat1* DSB formation.

### *h*^*+N*^ cells rarely generate *mat1* DSBs and typically carry two P alleles at *mat1*

Having established that HR is essential in *h*^*-S*^, we next asked why HR is dispensable in *h*^*+N*^. As we observed for suppressed *h*^*-S*^ mutants, the most probable explanation is inefficient DSB formation at the *mat1* locus in *h*^*+N*^. Indeed, southern blots and sequencing of Rad52-bound ssDNA previously revealed no detectable DSBs in *h*^*+N*^ (10,12). However, those reads were aligned to the *h*^*-S*^ reference genome, which differs substantially from *h*^*+N*^ at the *mat* locus (Figure 1C; (10,12)). Notably, the precise sequence of the *h*^*+N*^ *mat1* locus has, to our knowledge, not been published. To resolve this, we amplified the *h*^*+N*^ *mat1* region by PCR, yielding a ∼14 kb product that was cloned and sequenced. As expected, the sequence contained the duplicated *mat2*-K-*mat3* region (File S2). Comparison with the PomBase *h*^*90*^ reference revealed 36 differences within the K region. These sequence discrepancies were consistently present in our *h*^*+N*^ and *h*^*90*^ strains, in Bioneer library strains, and in other sequenced *S. pombe* isolates (34). Thus, the PomBase *h*^*90*^ K-region reference sequence contains 36 sequencing errors that should be corrected (15).

Unexpectedly, in contrast to the *mat2-P* and *mat3-M* arrangement in *h*^*90*^, our *h*^*+N*^ clones consistently carried P alleles on both sides of *mat1* (Figure 3A). To quantify the allelic composition flanking *mat1*, we performed qPCR on genomic DNA using primer pairs with one primer annealing to the sequence adjacent to *mat1* (Figure 3B, orange arrows) and one allele-specific primer (violet arrows). As controls, *h*^*90*^ contained both M and P alleles in roughly equal proportions, whereas *h*^*-S*^ carried only the M allele. The apparent two-fold higher M signal in *h*^*-S*^ reflects that only half of *h*^*90*^ cells harbor *mat1-M*, with the other half carrying *mat1-P*. Consistent with our previous results, *h*^*+N*^ clones contained exclusively the P allele on the left-hand side of *mat1*, while the right-hand side also predominantly carried the P allele, approximately 20-fold more frequently than M (Figure 3B, right panel). These results confirm that *h*^*+N*^ typically harbors two P alleles at *mat1*.

**Figure 3.**
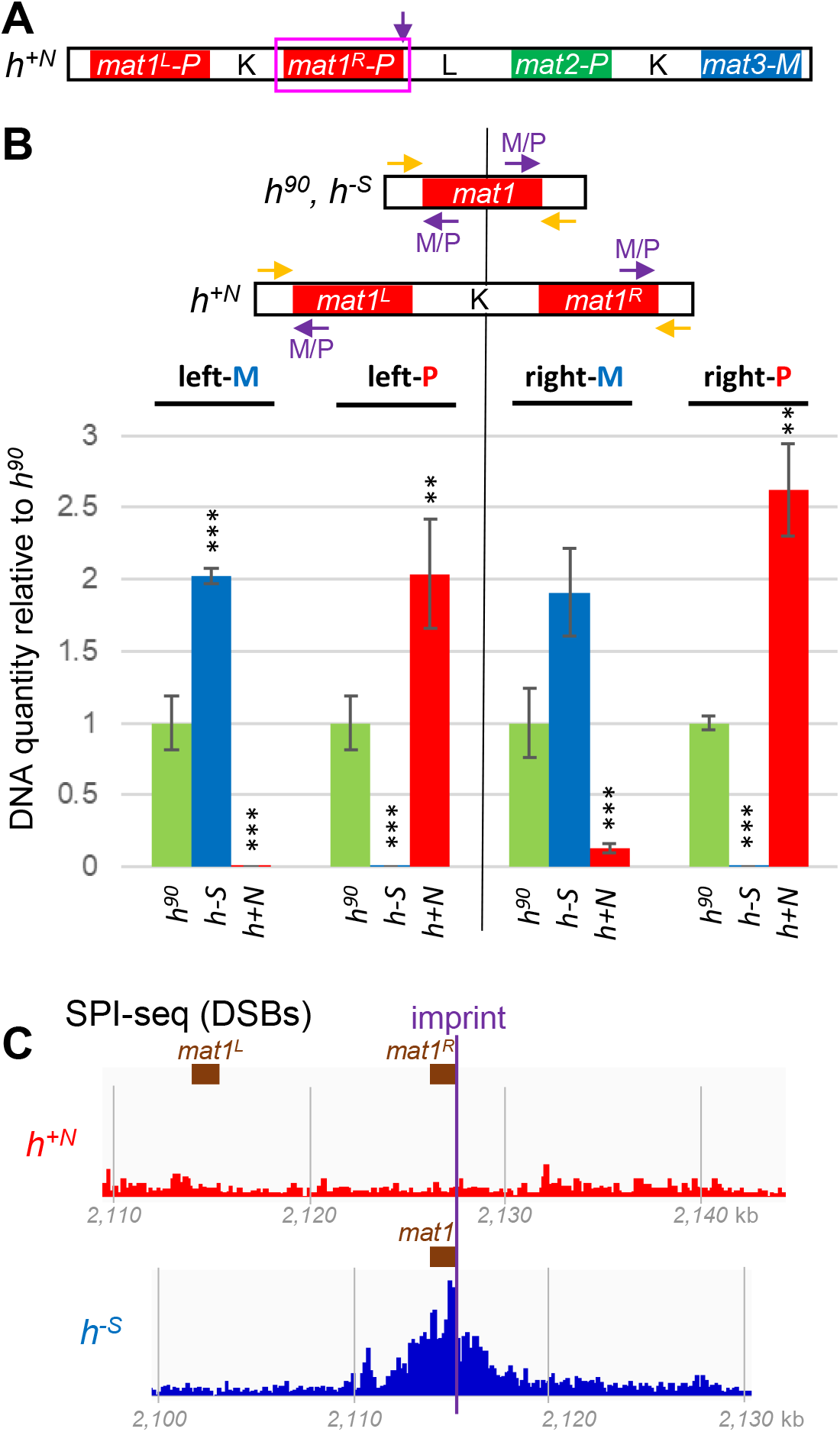
*h*^*+N*^ typically carries P alleles on both sides of *mat1* and lacks detectable *mat1* DSBs. (**A**) Schematic representation of the *mat* locus organization in *h*^*+N*^ showing P alleles on both sides of *mat1*. (**B**) Quantification of P and M alleles at both sides of *mat1* by qPCR. Genomic DNA was amplified using primer pairs consisting of one primer annealing to the sequence adjacent to *mat1* (orange arrows) and one allele-specific primer (violet arrows; Table S3). In *h*^*90*^ and *h*^*–S*^ strains, which harbor only a single cassette at *mat1*, left- and right-side qPCRs yield identical results. Data are expressed as DNA levels relative to *h*^*90*^ after normalization to *act1*. Bars represent the mean of three independent biological replicates ± standard deviation (SD). Asterisks indicate statistical significance (** P < 0.01; *** P < 0.001, two-tailed Student’s t-test) when comparing heterothallic strains with *h*^*90*^. (**C**) *mat1* DSBs are absent in most *h*^*+N*^ cells. ssDNA-associated Rad52 was immunoprecipitated, and the recovered DNA was sequenced using SPI-seq (12). Reads from the published *h*^*+N*^ dataset were aligned to our *h*^*+N*^ sequence, and reads from the *h*^*-S*^ dataset to the *h*^*-S*^ reference sequence, then plotted across the *mat1* locus. Coordinates (kb) are given relative to the respective reference sequences.

To assess whether *h*^*+N*^ contains additional genomic changes, we performed Illumina sequencing and detected no major differences relative to the *h*^*-S*^ reference outside of the *mat* locus (File S1). Using our *h*^*+N*^ sequence (Files S3 and S4), we then reanalyzed the Rad52-bound ssDNA dataset from Zhou et al. (12). Unlike in *h*^*-S*^, no enrichment was observed around *mat1* in *h*^*+N*^, confirming that DSBs are absent in most *h*^*+N*^ cells (Figure 3C).

### The *mat1* locus of *h*^*+N*^ nucleates H3K9me, leading to transcriptional silencing

We next investigated why DSB formation is inefficient at the *mat* locus of *h*^*+N*^. The absence of detectable changes in the *h*^*+N*^ DNA sequence around the *mat1* imprint site, or of potential suppressor mutations when compared to *h*^*-S*^, suggested that the reduction in DSB production is epigenetically regulated. Because *mat1* in *h*^*+N*^ carries a duplication of the K region, it also harbors the *cenH* and *REIII* elements that promote heterochromatin assembly (Figure 1). We therefore hypothesized that these elements nucleate heterochromatin at *mat1* in *h*^*+N*^, leading to transcriptional silencing and inhibition of DSB formation.

To test this, we examined heterochromatin status around *mat1* in *h*^*+N*^ using H3K9me2 ChIP-qPCR. Due to sequence identity between the *mat1* duplication and *mat2*-K-*mat3*, most of *mat1* cannot be uniquely assayed; we therefore focused on flanking unique regions, including the imprint site. Importantly, H3K9me2 levels were strongly increased around the imprint site in *h*^*+N*^ compared to *h*^*90*^, *h*^*-S*^, or *h*^*+N*^ *clr4Δ* lacking the sole H3K9 methyltransferase in *S. pombe* (Figure 4A). H3K9me2 enrichment gradually declined across the L region toward *mat2-P*. Notably, methylation levels around the imprint site in *h*^*+N*^ were comparable to those at the silent *mat3-M* locus (Figure 4A). In *h*^*-S*^, the absence of the K region resulted in reduced *mat3-M* methylation. Reanalysis of published H3K9me2 ChIP-seq datasets confirmed the qPCR results: *h*^*+N*^ displayed strong H3K9me2 enrichment around both *mat1* and *mat2/3*, whereas only *mat2/3* was methylated in *h*^*90*^ and *h*^*-S*^ (Figures 4B and S1; (35–39)).

**Figure 4.**
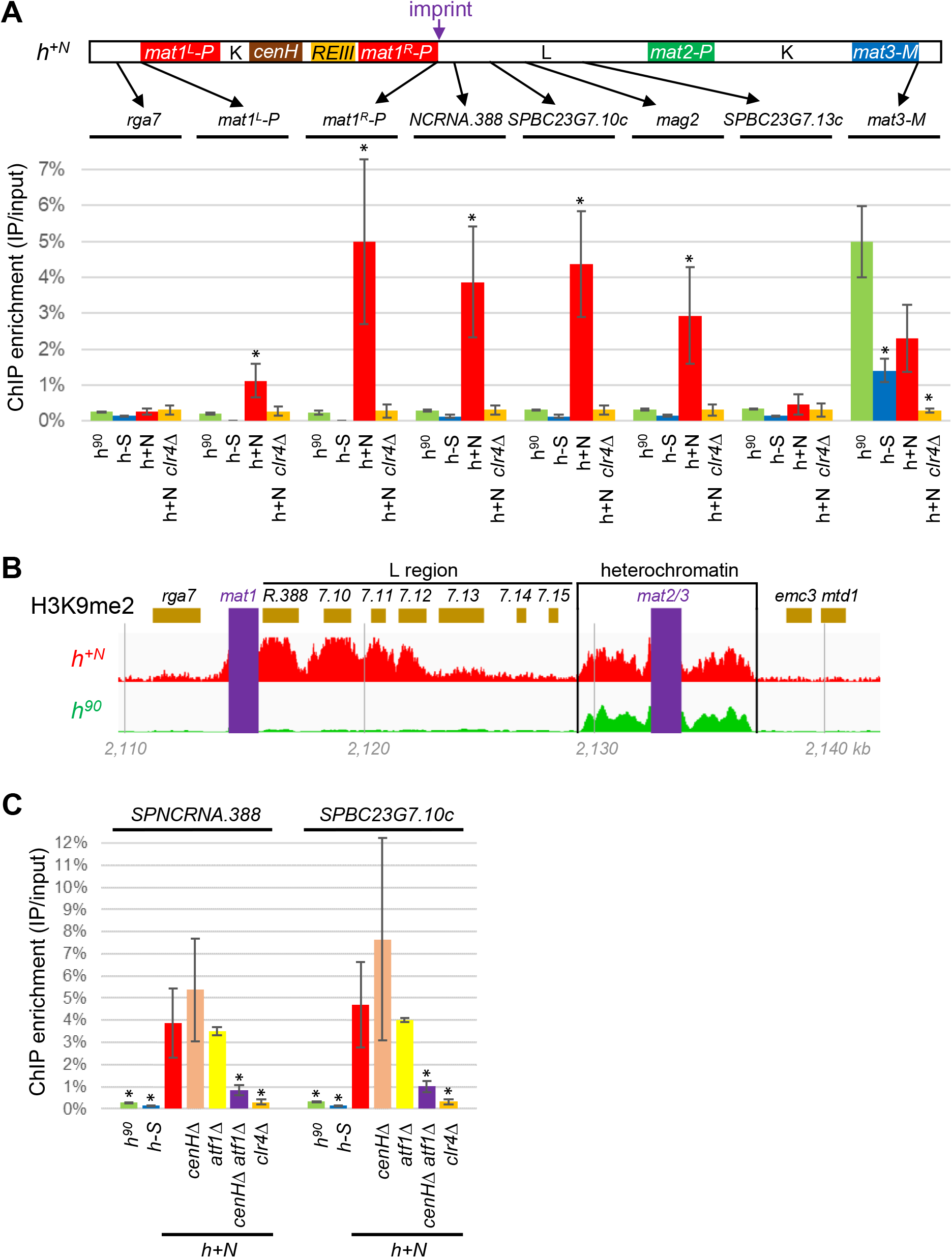
H3K9me2 enrichment around the *mat1* imprint site is strongly increased in the *h*^*+N*^ strain and requires *cenH* and Atf1. (**A**) Quantification of H3K9me2 occupancy by ChIP-qPCR. Data are shown as the percentage of immunoprecipitated DNA (IP) relative to total input DNA. Values represent the mean of two independent biological replicates ± SD; asterisks indicate P < 0.05 (two-tailed Student’s t-test) when comparing each strain with *h*^*90*^. (**B**) H3K9me2 enrichment determined by ChIP-seq. Published datasets for *h*^*+N*^ (35) and *h*^*90*^ (38) were plotted across the *mat* locus, with repetitive sequences (*mat1, mat2, mat3*, and the K region) omitted and indicated by violet rectangles. Gene names above the track are abbreviated: *SPNCRNA*.*388, SPBC23G7*.*10c, SPBC23G7*.*11* (*mag2*), *SPBC23G7*.*12c* (*rpt6*), *SPBC23G7*.*13c, SPBC23G7*.*14*, and *SPBC23G7*.*15c* (*rpp202*). Coordinates (kb) are given relative to the *h*^*-S*^ reference sequence. (**C**) Dependence of H3K9me2 enrichment on *cenH* and Atf1. Occupancy in the indicated strains and loci was quantified as described in (A), except that asterisks denote statistical comparison to *h*^*+N*^.

To determine whether this heterochromatin depends on *cenH* and the Atf1-bound *REIII* element, we analyzed the effect of their deletion. Single deletions of *cenH* or *atf1* did not substantially reduce H3K9me2, but the double mutant showed a strong loss of methylation (Figure 4C), similar to previous findings at the *mat2/3* locus (4).

Since H3K9me typically induces transcriptional silencing, we next examined expression of genes near the imprint site. Compared to *h*^*90*^ or *h*^*-S*^, the *h*^*+N*^ strain exhibited markedly reduced transcription of *SPNCRNA*.*388, SPBC23G7*.*10c*, and *mag2* (Figure 5A). Expression of *mat1-Pc* in *h*^*+N*^ was similar to *h*^*90*^, whereas *mat1-Mc* was approximately twofold higher in *h*^*-S*^ than in *h*^*90*^. This increase is expected, as in *h*^*90*^ only half of the cells express *mat1-Mc* while the other half express *mat1-Pc*. Together, these results suggest partial silencing of the left-hand *mat1-Pc* allele in *h*^*+N*^, consistent with the elevated H3K9me2 levels observed at this locus (Figure 4A). As with H3K9me2, silencing was abolished in *clr4Δ* and strongly reduced in the *cenHΔ atf1Δ* double mutant (Figure 5B). Unlike H3K9me2 levels, however, transcription was also affected in the *cenHΔ* and *atf1Δ* single mutants, indicating that gene expression is a more sensitive measure of heterochromatin defects. Together, these findings demonstrate that duplication of *cenH* and *REIII* to the *mat1* locus in *h*^*+N*^ nucleates heterochromatin formation, characterized by H3K9me and transcriptional silencing of adjacent genes.

**Figure 5.**
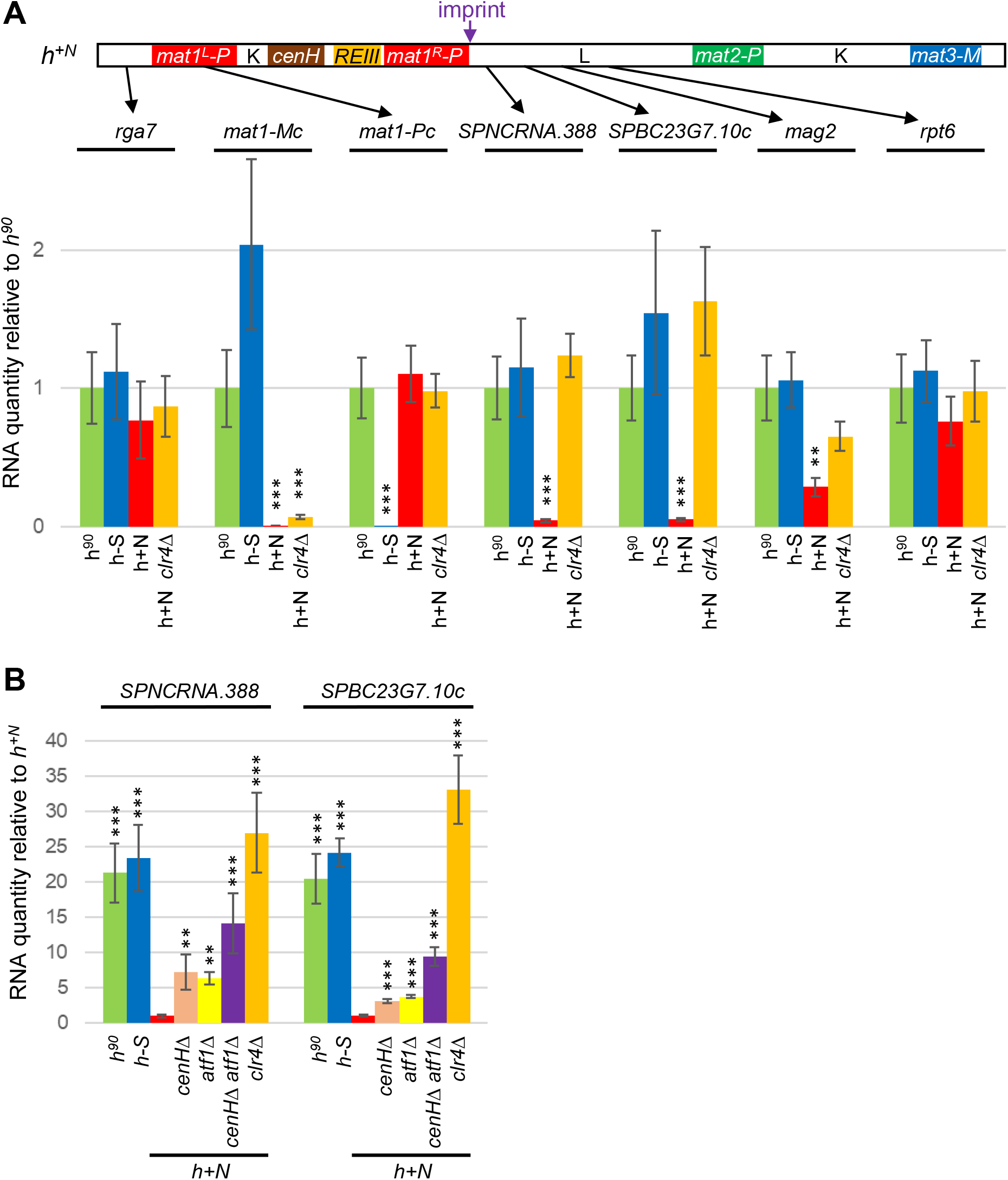
Transcription at *mat1* and adjacent genes is silenced in *h*^*+N*^. (**A**) Gene expression around the *mat1* locus. RNA was isolated from exponentially growing cells (OD600 ≈ 0.4) and analyzed by RT-qPCR. Transcript levels are shown relative to *h*^*90*^, normalized to *act1*. Values represent the mean ± SD of two independent biological replicates. Statistical significance was assessed using two-tailed Student’s t-tests, comparing each strain to *h*^*90*^ (**P < 0.01, ***P < 0.001). (**B**) Gene expression was analyzed as in (A), except transcript levels are shown relative to *h*^*+N*^. Values represent the mean ± SD of three independent biological replicates.

### Disruption of heterochromatin restores switching, HR essentiality, and imprintosome recruitment

If our hypothesis that the *mat1* heterochromatin in *h*^*+N*^ prevents DSB formation is correct, then inhibition of H3K9me should reactivate switching at the right-hand *mat1* allele, which is predominantly of the P type (Figure 3B). Consistent with this prediction, deletion of *clr4* almost completely converted the right-hand allele to M, while the left-hand allele largely remained P (Figures 6A and S2). Similarly, reversion from the *mat1* P–P configuration to P–M was also detected in *atf1*Δ, *clr6-1, dcr1*Δ, and *cenH*Δ mutants, all of which disrupt maintenance of the silent *mat2/3* locus (Figure 6A).

**Figure 6.**
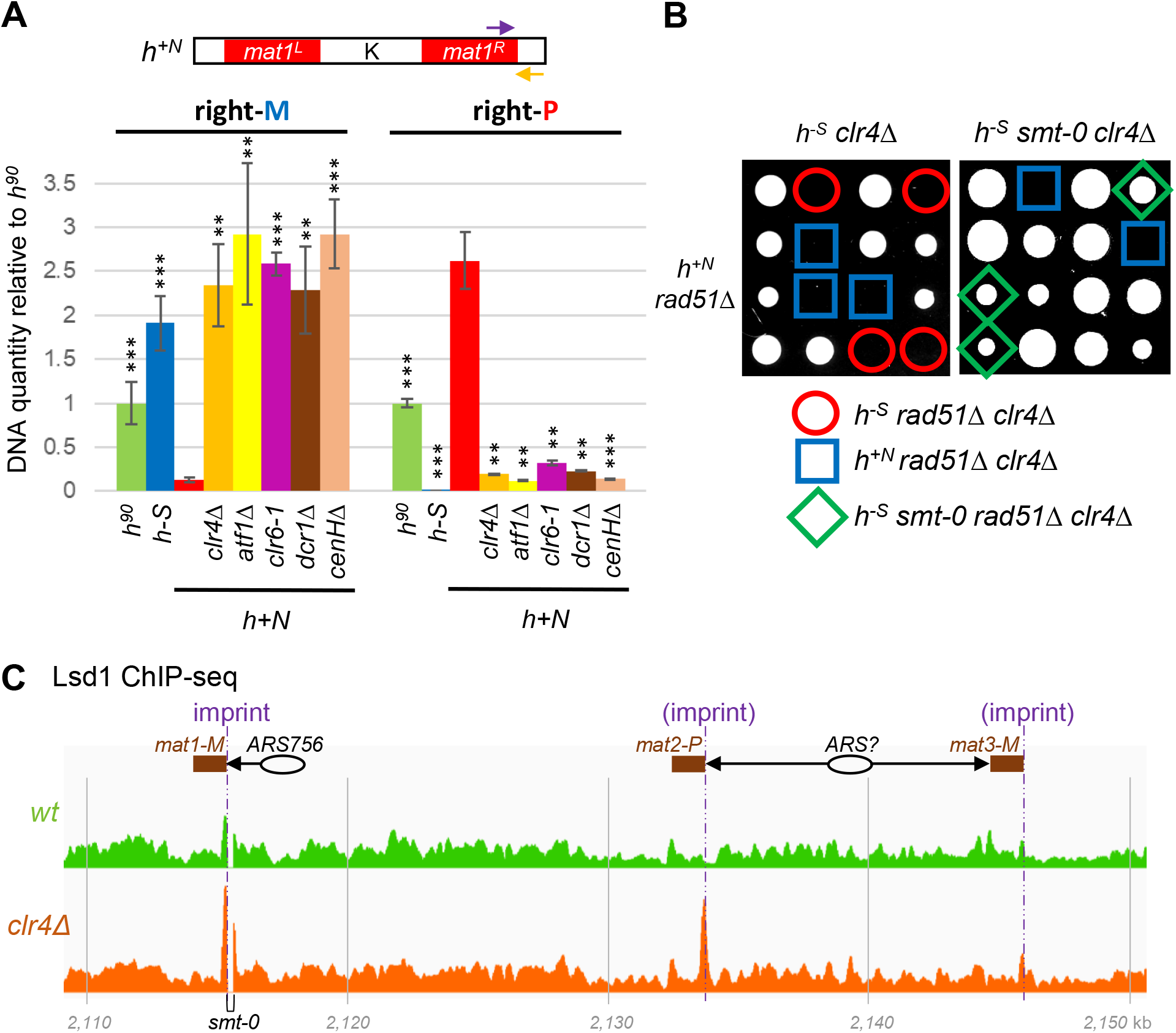
Disruption of heterochromatin triggers switching, HR essentiality, and imprintosome recruitment. (**A**) Mutations affecting heterochromatin formation cause the right-hand *mat1* allele in *h*^*+N*^ cells to switch to the M state. The relative abundance of P and M alleles at the right-hand side of *mat1* was quantified by qPCR. DNA levels were normalized to *act1* and expressed relative to *h*^*90*^. Data are shown as mean ± SD from two independent biological replicates. Statistical significance was determined using two-tailed Student’s t-tests, comparing each strain to *h*^*+N*^ (**P < 0.01, ***P < 0.001). (**B**) Loss of H3K9me causes synthetic lethality with *rad51Δ* in *h*^*+N*^ cells. The indicated strains were crossed, and meiotic progeny were analyzed by tetrad dissection. (**C**) Lsd1 enrichment at *mat2-P* is elevated in the absence of H3K9me. Lsd1-FTP enrichment was assessed by ChIP-seq (42). Published datasets from wild-type (*mat1-Msmt-0 Lsd1-FTP::NatN2*) and *clr4Δ* strains are shown across the *mat* locus. Sequences identical to the imprint site at *mat1-M* and the predicted directions of replication are indicated. Ellipses mark origins of replication (ARS, autonomously replicating sequence). Because the *mat1-M* and *mat3-M* sequences are identical, enrichment in this region cannot be uniquely assigned to either locus. Coordinates (kb) are relative to our *h*^*90*^ *mat1-M* reference genome sequence (Files S5 and S6).

We next tested whether heterochromatin removal in *h*^*+N*^ also reinstates the requirement for HR for survival. As expected, the *rad51*Δ *clr4*Δ double mutant was lethal not only in *h*^*-S*^ cells but also in *h*^*+N*^ (Figure 6B). By contrast, the *h*^*-S*^ *smt-0 rad51*Δ *clr4*Δ strain was viable, confirming that lethality is specifically caused by *mat1* DSBs. Together, these results demonstrate that heterochromatin at the *mat1* imprint site in *h*^*+N*^ suppresses DSB formation, allowing survival without HR. When heterochromatin is disrupted, DSBs are restored and HR once again becomes essential for viability.

But how does heterochromatin block *mat1* DSB formation? We hypothesize that it does so by preventing the binding of the imprintosome complex, composed of Lsd1, Lsd2, Phf1, and Phf2, whose Pfh1/2 subunits bind to histone marks according to their methylation status (1). This idea is supported by the absence of both imprinting and imprintosome association at the silent *mat2/3* locus, even though the sequences at the centromere-distal ends of the M and P alleles required for imprinting are present (40,41). To determine whether H3K9me influences imprintosome recruitment, we analyzed published Lsd1 ChIP-seq datasets from the *mat1-Msmt-0* strain (an *h*^*90*^ derivative with stable *mat1-M*) and its *clr4Δ* mutant (42), using our *h*^*90*^ reference genome (Files S5 and S6). Strikingly, loss of H3K9me markedly increased Lsd1 binding at the imprint sequence of *mat2-P*, reaching levels comparable to those observed at *mat1-M* (Figure 6C). By contrast, only a modest increase was detected at the unique sequence downstream of *mat3-M*. This difference is consistent with the expected directions of replication, as the K region likely contains an early origin of replication. Accordingly, both *mat2-P* and *mat1-M* are replicated in the same direction, where replication fork pausing, and thus imprinting, is expected to occur, whereas *mat3-M* is replicated in the opposite direction ((40,43); Figure 6C). Together, these results demonstrate that H3K9me-dependent heterochromatin blocks imprintosome recruitment and thereby prevents *mat1* DSB formation.

## DISCUSSION

In this study, we highlight a key difference in programmed DSB formation between the two heterothallic laboratory strains most widely used in research. While *mat1* DSBs are generated efficiently in *h*^*-S*^, they are nearly absent in *h*^*+N*^. We demonstrate that this absence in *h*^*+N*^ results from the duplication of the K region into the *mat1* locus, which triggers heterochromatin nucleation. The resulting heterochromatin spreads to the imprint site, preventing imprintosome recruitment and silencing nearby genes. The most striking phenotypic consequence of this difference in DSB formation is the lethality of HR mutants in *h*^*-S*^, contrasted with their viability in *h*^*+N*^ (Figures 1 and 7).

**Figure 7.**
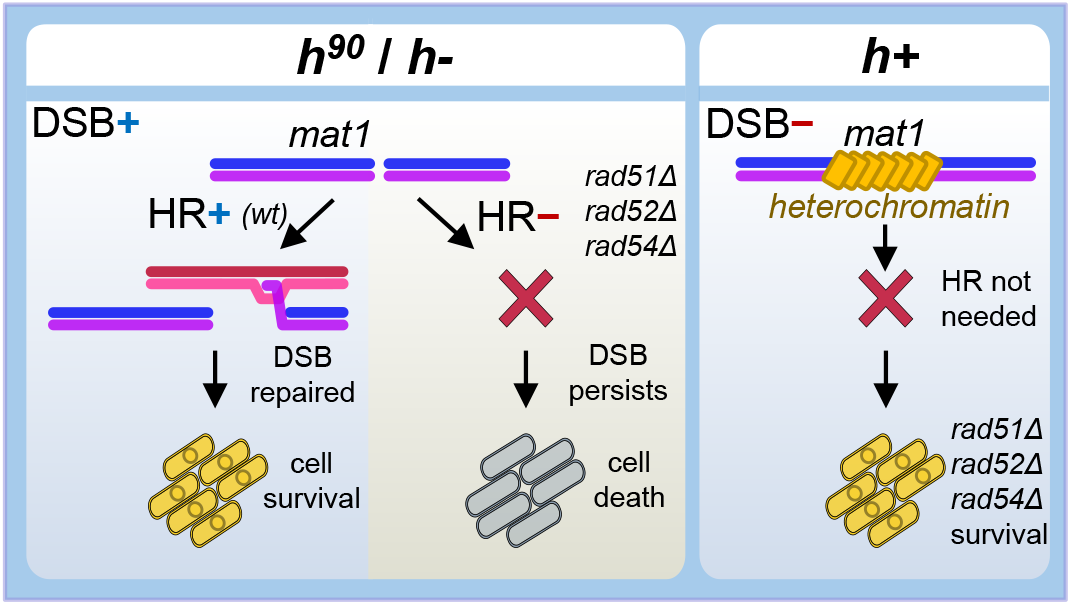
Graphical summary. Schematic overview of differences in HR essentiality among commonly used *S. pombe* strains. Left panel: In the *h*^*90*^ and *h*^*-S*^ strains, a programmed DSB is efficiently generated at the *mat1* locus and repaired by HR under wild-type conditions. Middle panel: When HR is defective, *mat1* DSBs remain unrepaired, leading to cell death. Right panel: In the *h*^*+N*^ strain, the *mat1* imprint site is embedded within heterochromatin, which prevents DSB formation. Consequently, HR is not required for survival, and HR mutants remain viable.

### *h*^*-S*^: *mat1* DSBs enforce HR essentiality unless their generation is suppressed

The observed indispensability of HR for the survival of *h*^*-S*^ strains is not unexpected, given that *h*^*-S*^ generates *mat1* DSBs with an efficiency comparable to *h*^*90*^ (10,12). Note that the DSB signals observed in early Southern blot analyses (e.g., (10)) likely represent imprinting events rather than bona fide breaks (44). Nevertheless, because these imprints ultimately give rise to DSBs, we refer to them as DSBs here for conceptual simplicity. Surprisingly, many *S. pombe* researchers appear unaware of the essential role of HR in *h*^*-S*^ and continue to employ *rad51Δ, rad52Δ*, or *rad54Δ* strains in the *h*^*-S*^ background (Table S1). This misconception can be traced to the study by Ostermann et al., which reported that Rad52 is essential for viability in *h*^*90*^, but dispensable in *h*^*+N*^ and *h*^*-S*^ heterothallic strains (19). Since the tetrad analysis supporting this conclusion was never published, it is plausible that their assessment relied on limited or incomplete data. The misunderstanding persists today in PomBase, which states that deletions of *rad51, rad52*, and *rad54* are viable, but without specifying the genetic background (15). While this statement is accurate for *h*^*+N*^ strains, it does not hold true for the wild-type *mat1* contexts of *h*^*90*^ and *h*^*-S*^.

After confirming that major HR mutants are lethal in *h*^*-S*^ (Figure 1D), we examined publicly available *h*^*-S*^ *rad51Δ* strains. We found that they were either mislabeled (incorrectly annotated as *h-* or *rad51Δ*) or harbored additional mutations likely suppressing *mat1* DSB formation. These probable cases of mislabeling or hidden suppressor mutations in published *rad51Δ, rad52Δ*, and *rad54Δ h-* strains raise concerns about the reliability of the conclusions drawn from such strains (Table S1). We demonstrated that suppressors might include *smt-0, swi1Δ*, or *fml1Δ* (Figure 2).

Interestingly, despite extensive research on proteins required for MTS, a role for Fml1 in this process has not been previously recognized (Figure 2D). Its absence from high-throughput MTS screens (45) is likely explained by the wild-type *fml1* allele that we observed in the strain annotated as *fml1Δ* in the Bioneer library. Although Fml1’s precise contribution to MTS remains to be elucidated, its known activities in replication fork reversal and in the restoration of regressed forks suggest plausible roles at the imprint site, where chicken-foot structures have been observed (46–48).

### *h*^*+N*^: *mat1* DSBs are suppressed by heterochromatin formation

Although the deficiency in *mat1* DSB generation in *h*^*+N*^ was first documented more than four decades ago (10), its underlying cause has not been directly addressed, and its broader implications for work with heterothallic strains remain underappreciated. Notably, when *h*^*+N*^ isolates are freshly derived from *h*^*90*^, their *mat1* DSB frequency is only about fourfold lower than in *h*^*90*^ (10). This is expected, as establishment of *mat2/3* heterochromatin can require up to 20 generations (49), and heterochromatin assembly at *mat1* would likely be considerably slower due to the absence of the *REII* element and inverted repeats (Figure 1).

Fresh *h*^*+N*^ isolates are also expected to carry the M allele (a copy of *mat3-M*) on the right-hand side of *mat1*, yet we found that most cells in our *h*^*+N*^ strain instead carry the P allele (Figure 3). This agrees with the digestion patterns reported by Beach and Klar, although the absence of the M allele at this position was not noted (10). Why then did the early *h*^*+N*^ strain with a *mat1-P-M* configuration predominantly evolve into *P-P*? We propose that simultaneous expression of both P and M information is disadvantageous, and cells with a P-P configuration and a consistent P phenotype gradually outcompeted their ambiguous siblings. A similar phenomenon is observed when heterothallic cells progressively dominate homothallic cells under laboratory conditions (1).

Despite clear evidence of H3K9me2 spreading from *h*^*+N*^ *mat1* in published ChIP-seq datasets, this phenomenon has not been noted in the corresponding studies (35–37,39). Our hypothesis that heterochromatin at *mat1* arises from the combined action of *cenH* and *REIII* is consistent with the work of Wang et al., who showed that recruitment of Clr4 together with REIII can establish silencing even in the absence of inverted repeats and REII (50). Although the effect they reported was weaker than what we observe in *h*^*+N*^, their experiment lasted only four days, whereas currently used *h*^*+N*^ strains have been propagated for decades. Moreover, the proposed selective pressure to avoid simultaneous P and M expression may have also contributed to heterochromatinization and silencing of the right-hand *mat1* allele. This right-hand allele is completely silent (10), explaining why *h*^*+N*^ expresses only P information from the left-hand allele.

We support our hypothesis that *mat1* heterochromatin in *h*^*+N*^ prevents DSB formation based on two observations involving heterochromatin mutants: they enable the restart of MTS at the right-hand allele, and deletion of *clr4* restores HR essentiality (Figure 6). This hypothesis is further supported by the findings of Lorentz et al., who showed that deletion of *swi6* in *h*^*+N*^ induced DSBs at the right-hand *mat1* allele (51). The near-complete conversion of this allele to M after switching reactivation (Figure 6A) likely reflects the inherent bias in switching toward the M donor allele in both cells expressing the P allele (as in *h*^*+N*^) and heterochromatin mutants (1).

We further demonstrate that H3K9me inhibits DSB formation by preventing recruitment of the imprintosome complex (Figure 6C). This finding provides a direct molecular explanation for how heterochromatin suppresses *mat1* cleavage. Note that Liu et al. did not observe the pronounced Lsd1/2 enrichment at the *mat2-P* locus in *clr4Δ* cells because they used the *h*^*–S*^ strain as a reference, which carries a *mat2-P* deletion (42). Our findings, however, contrast with those of Raimondi et al., who reported that deletion of *clr4* does not promote Lsd1/2 recruitment at *mat2/3* (41). However, as those data were not shown, their conclusions cannot be independently verified. Notably, even though the imprintosome can bind to *mat2-P* in the absence of H3K9me, the lack of the downstream sequence containing Sap1 binding sites would still prevent imprint stabilization and DSB formation at this locus. A similar situation likely occurs at the left-hand *mat1-P* allele in *h*^*+N*^ cells, where the low level of H3K9me (Figure 4A) may permit imprintosome binding and trigger the replication fork pausing previously observed (43).

### Is HR essential

Various organisms, including *S. pombe, S. cerevisiae*, and vertebrates, share a common feature: HR is essential for the survival of some cells but dispensable in others. Here, we show that in *S. pombe*, cells lacking HR are inviable unless *mat1* DSB production is suppressed, for example, through mutations such as *smt-0, swi1Δ, fml1Δ*, or the *mat2/3* duplication in *h*^*+N*^.

A similar situation occurs in *S. cerevisiae*: “wild-type” homothallic strains expressing the HO endonuclease, which produces DSBs during MTS, are inviable in the absence of HR (14). In contrast, most laboratory *S. cerevisiae* strains are heterothallic (stably *MAT*a or *MAT*α) due to the absence of HO endonuclease. In these strains, the lack of *MAT* DSBs renders HR mutants viable (52,53).

In vertebrates, HR mutants are generally viable in differentiated cells, but HR is particularly critical in rapidly proliferating cells, such as embryonic stem cells. This likely underlies the embryonic lethality observed in key HR mutants (54–58).

Thus, should HR be considered essential or dispensable for survival in these organisms? We propose that these are not mutually exclusive statements: HR essentiality depends on specific cellular contexts, and the conditions under which it is evaluated must be carefully considered.

### Suggestions for working with strains of different mating types

Both widely used heterothallic strains arose from rearrangements of the *mat* locus, which render them unstable over long-term use. *h*^*+N*^ strains tend to further expand the *mat* locus over time, increasing the number of *mat* cassettes and intervening regions to multiple copies ((10); our observations). *h*^*-S*^ cells, on the other hand, can undergo recombination between the *mat1-M* and *mat3-M* alleles, resulting in deletion of the essential L region along with *mat3-M* from chromosome II and its subsequent propagation as a plasmid (10). Based on these observations and the differences between *h*^*-S*^ and *h*^*+N*^ discussed above, we propose the following guidelines:

1, Determine mating type for every strain. When evaluating the effect of a mutation, both the wild-type and mutant strains must share the same background. Surprisingly, this seemingly obvious rule is often neglected.

2, Avoid using HR mutants in *h*^*-S*^ or *h*^*90*^ backgrounds when studying MTS-independent roles of HR. Such mutants may be evaluated in *h*^*+N*^, unless combined with mutations that disrupt *mat* heterochromatin (e.g., *clr4Δ*, Figure 6B).

3, Ideally abolish *mat1* DSBs for studies not evaluating MTS. For example, using the *mat1-Msmt-0* and *mat1-PΔ17* deletions is recommended (21,22,59). This approach is analogous to most *S. cerevisiae* laboratories, which utilize strains lacking the HO endonuclease to prevent programmed DSB formation.

4, Freeze all newly generated mutants immediately and sequence their genomes. Genome sequencing is crucial to exclude suppressor mutations, which are often present in published strains and can confound experimental conclusions. Illumina sequencing of an *S. pombe* strain, including library preparation, can currently be performed for less than $100.

## MATERIALS AND METHODS

### Yeast strains

*Schizosaccharomyces pombe* strains were cultured following standard protocols in yeast extract medium with supplements (YES; (60)). Genetic manipulations were performed according to established procedures (61). The genotypes of all strains used in this study are listed in Table S2.

The strain carrying a 4.3 kb *cenH* deletion at the *h*^*+N*^ *mat1* locus (yPK799-10) was generated by transforming strain yPK795-8B (derived from the Bioneer library base strain) with a DNA fragment in which *cenH* was replaced by *ura4*. The transformation cassette was obtained by digesting plasmid pPK415 with EcoRI and SacI. Transformants were selected on uracil-deficient plates, and correct integration was confirmed by DNA sequencing.

For tetrad analysis, strains were crossed and sporulated on EMM-N plates at 28 °C for 2–3 days. Tetrads were dissected using a Singer MSM300 dissection microscope (Singer Instruments), and individual spores were incubated on YES plates at 28 °C for 5–7 days.

### Quantification of genomic DNA

Cells were grown in 2 ml YES medium to an optical density (OD600) of approximately 1, and genomic DNA was extracted using the Dr. GenTLE (from Yeast) High Recovery Kit (Takara).

qPCR reactions (20 μl total volume) contained 25 ng of genomic DNA, 4 pmol of each primer (Table S3), and 1× HOT FIREPol EvaGreen qPCR Mix Plus (Solis BioDyne). Amplification was performed on a StepOnePlus Real-Time PCR System (Applied Biosystems) with the following cycling conditions: 12 min at 95 °C; 40 cycles of 95 °C for 15 s, 60 °C for 20 s, and 72 °C for 20 s. Signal normalization was carried out using the *act1* reference gene.

### H3K9me2 ChIP-qPCR

Exponentially growing cells (OD600 ≈ 0.5) in YES medium were crosslinked with 1% formaldehyde for 10 min at room temperature (RT). Crosslinking was quenched with 125 mM glycine for 10 min. Cells were washed with PBS and lysed with glass beads.

Chromatin was sheared using a Bioruptor sonicator (Diagenode) for 20 cycles (30 s ON / 30 s OFF, high power). For each immunoprecipitation (IP), 4.5 mg of total protein was used; 3% of the chromatin extract was reserved as input control. IPs were performed with 3 μg of anti-H3K9me2 antibody (ab1220, Abcam) and incubated for 2 h at 4 °C with rotation. Subsequently, 50 μl of BSA-blocked Dynabeads Protein G magnetic beads (Thermo Fisher) were added and incubated overnight at 4 °C with rotation.

The precipitated material and input were de-crosslinked and treated with RNase A and Proteinase K. DNA was purified using the MinElute PCR Purification Kit (Qiagen) and eluted into 10 μl of water. DNA enrichment was quantified by qPCR as described above, using 5 μl of 100x diluted DNA and primers listed in Table S3. Enrichment was calculated as: %IP/Input = 2^[Ct(Input)−Ct(IP)] × 3%.

### Quantification of RNA

Cells were cultured in 10 ml YES medium to OD600 ≈ 0.4, harvested by centrifugation (1,000 × g, 1 min), washed in 1 ml sterile water, transferred to 2 ml screw-cap tubes, pelleted again, and flash-frozen in liquid nitrogen. Cell lysis was performed with 0.5 mm acid-washed glass beads (Sigma-Aldrich) and 0.5 ml TRI Reagent (MRC) using a FastPrep-24 grinder (MP Biomedicals) at 6.5 m/s for 1 min, followed by cooling on ice. Fifty microliters of BCP (1-bromo-3-chloropropane) were added, vortexed for 15 s, and incubated for 2 min at RT. Samples were centrifuged at 12,000 × g for 15 min at 4 °C, and 200 μl of the upper aqueous phase was transferred to a new tube. After an additional centrifugation (8 min, 12,000 × g, 4 °C), 150 μl of the supernatant was mixed with an equal volume of isopropanol to precipitate RNA. Samples were incubated for 5 min at RT and centrifuged (12,000 × g, 8 min, 4 °C). The pellets were washed with 1 ml of 75% ethanol, scraped from the tube walls, centrifuged (12,000 × g, 5 min, 4 °C), air-dried for 5 min, and dissolved in 100 μl DEPC-treated water.

For reverse transcription, 0.5 μg RNA was mixed with 0.5 μl of 100 μM oligo(dT)20 primer, 0.5 μl of 100 μM random hexamers (Generi Biotech), 1 μl of 10 mM dNTP mix (NEB), and water to 8.3 μl total. The mixture was denatured for 5 min at 65 °C, cooled on ice, and supplemented with 0.5 μl M-MuLV Reverse Transcriptase (NEB), 1 μl 10× RT buffer, and 0.2 μl SUPERase-In RNase Inhibitor (Invitrogen). Samples were incubated for 5 min at RT, followed by reverse transcription for 1 h at 42 °C and enzyme inactivation for 20 min at 65 °C. The resulting cDNA was diluted 50× with water and stored at −20 °C.

cDNA quantification (5 μl) was performed by qPCR as described for genomic DNA using primers listed in Table S3.

### Bioinformatic analyses

Genomic DNA for whole-genome sequencing was isolated using the Dr. GenTLE (from Yeast) High Recovery Kit (Takara). Library preparation and sequencing (PE150 reads, ∼1.2 Gb per sample) were performed by Novogene. Raw reads were trimmed with fastp (v1.0.1; with parameters --n_base_limit 10, -- qualified_quality_phred 5, --unqualified_percent_limit 50) and mapped to the *S. pombe* reference genome (972 *h-*, ASM294v3, February 2024 release) using BWA-MEM (v0.7.19). PCR and optical duplicates were identified using GATK MarkDuplicates (v4.6.2.0) and excluded from further analysis. HaplotypeCaller (ploidy = 1, GVCF mode) was used to generate variant calling files. Individual genomic variant calls were produced with GenotypeGVCFs and variants were filtered using VariantFiltration with the following criteria QD < 2.0 || FS > 60.0 || MQ < 40.0. Variants were annotated using SnpEff (v5.2). Structural variants were detected using Delly (v1.3.3).

ChIP-seq datasets were retrieved from ENA/SRA repositories using the SRA Toolkit (v3.0.10) and trimmed with Trimmomatic (v0.39; parameters SLIDINGWINDOW:4:20 MINLEN:20). Reads were aligned to the *S. pombe* genome using Hisat2 (v2.2.1; parameters: --no-spliced-alignment -k 15). For H3K9me ChIP-seq, the 972 *h*^−*S*^ reference genome was used. For Lsd1 ChIP-seq, reads were aligned to a modified genome incorporating the *h*^*90*^ sequence at the *mat* locus with the M cassette at *mat1*, prepared using Reform (https://github.com/gencorefacility/reform). Genome coverage tracks were calculated using deepTools (v3.5.4), normalized by counts per million and visualized in the Integrative Genomics Viewer (62).

SPI-seq reads obtained from the *h*^*+N*^ strain were trimmed as above and aligned to the modified *S. pombe* reference genome in which the *h*^*+N*^ *mat* region was inserted using Reform. Reads from the *h*^*-S*^ strain were aligned to the standard *h*^*-S*^ genome. Downstream processing was identical to ChIP-seq analyses.

## Abbreviations

HR: homologous recombination;
DSB: double-strand break;
MTS: mating-type switching;
H3K9me: histone H3 lysine 9 methylation;
ChIP: chromatin immunoprecipitation;
qPCR: quantitative polymerase chain reaction

## AUTHOR CONTRIBUTIONS

PK conceived and designed the study, performed the experiments, and wrote the manuscript. SP conducted all bioinformatic analyses. BS carried out the ChIP-qPCR experiments. All authors contributed to manuscript revision and approved the final version for submission.

## ACKNOWLEDGEMENTS

We thank Silvia Bagelova-Polakova for providing yeast strains. Funding from the Czech Science Foundation (CSF project GA23-05284S) and Masaryk University (GAMU grant MUNI/R/1262/2022) is acknowledged. We also acknowledge the use of ChatGPT (GPT-5, OpenAI) for assistance with language editing.

## Data availability statement

All data generated or analyzed during this study are available within the manuscript, the supporting information files, or the referenced publications. The raw DNA sequencing data have been deposited in the European Nucleotide Archive (ENA) under the study accession number PRJEB99022.

## SUPPORTING INFORMATION

Additional supporting information may be found online in the Supporting Information section at the end of the article.

**Figure S1**. H3K9me2 occupancy around the *mat1* imprint site is strongly increased in *h*^*+N*^

**Figure S2**. Heterochromatin mutants predominantly retain the P allele at the left-hand *mat1* site

**Table S1**. Mating types of *rad51Δ, rad52Δ*, and *rad54Δ* strains used in recent publications

**Table S2**. *S. pombe* strains used in this study

**Table S3**. Primers used in this study

**File S1**. DNA variants identified by whole genome sequencing

**File S2**. Sequence of *mat* locus in the *h*^*+N*^ strain

**File S3**. Full *S. pombe h*^*+N*^ sequence (FASTA)

**File S4**. *S. pombe h*^*+N*^ sequence features (GFF3)

**File S5**. Full *S. pombe h*^*90*^ sequence containing the M cassette at *mat1* (FASTA)

**File S6**. *S. pombe h*^*90*^ sequence features (GFF3)

