## Supplementary figures and tables for "Homologous recombination mutant lethality differs between *h*− and *h+ Schizosaccharomyces pombe* strains due to *mat1* heterochromatin"

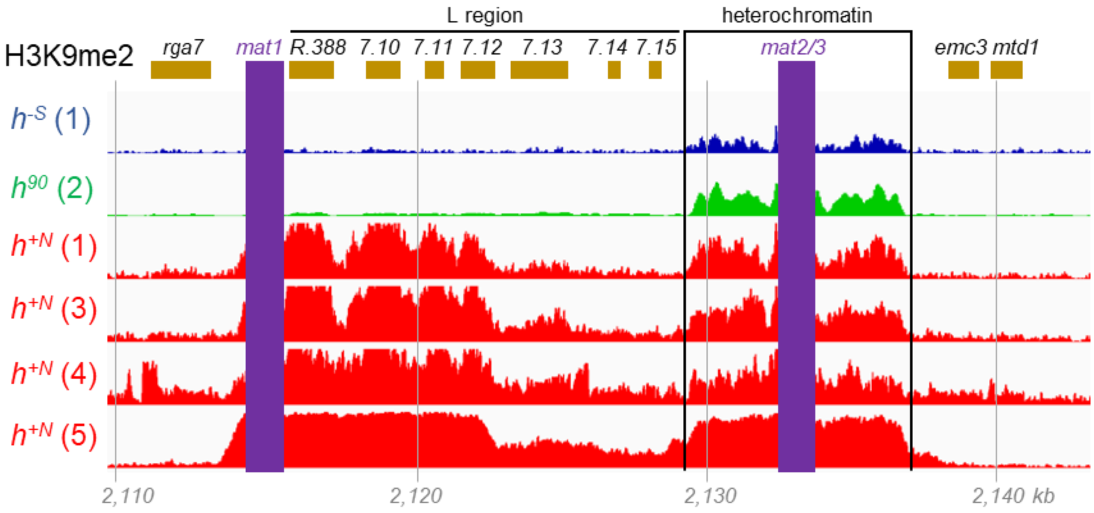


**Figure S1**. H3K9me2 occupancy around the *mat1* imprint site is strongly increased in $h^{+N}$. H3K9me2 enrichment was determined by ChIP-seq in previous studies, with the numbers in parentheses (1–5) corresponding to the reference list below. The datasets were plotted across the *mat* locus, with repetitive sequences (*mat1, mat2, mat3*, and the K region) omitted and indicated by violet rectangles. Gene names above the track are abbreviated: *SPNCRNA.388, SPBC23G7.10c, SPBC23G7.11* (*mag2*), *SPBC23G7.12c* (*rpt6*), *SPBC23G7.13c, SPBC23G7.14*, and *SPBC23G7.15c* (*rpp202*). Coordinates (kb) are given relative to the $h^{-S}$ reference sequence.

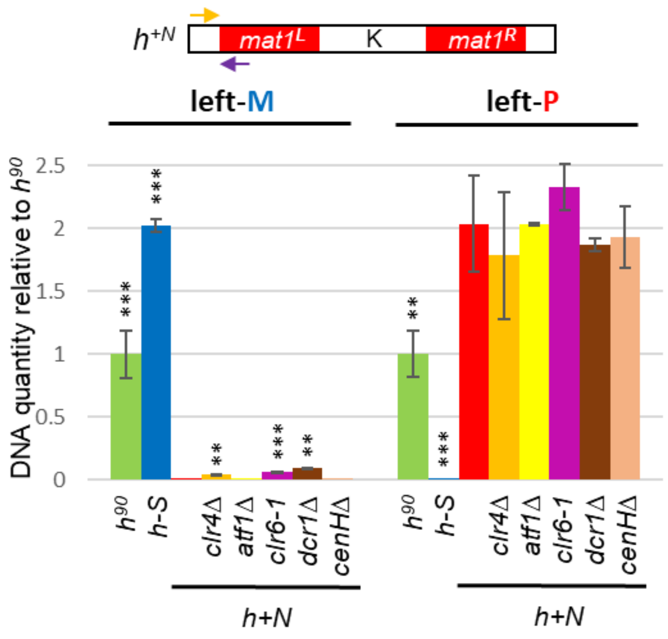


**Figure S2**. Heterochromatin mutants predominantly retain the P allele at the left-hand *mat1* site. The relative abundance of P and M alleles at the left-hand side of *mat1* was quantified by qPCR. DNA levels were normalized to *act1* and expressed relative to $h^{90}$. Data represent mean ± SD from two independent biological replicates. Statistical significance was assessed using two-tailed Student's t-tests, comparing each strain to $h^{+N}$ (**P < 0.01, ***P < 0.001).

### Supplementary tables

**Table S1**. Mating types of *rad51Δ*, *rad52Δ*, and *rad54Δ* strains used in recent publications. Studies containing the mutants in the *h-* background are highlighted red.

| year | First author | Last author | Pubmed ID | mating type used | problematic strains |
|---|---|---|---|---|---|
| 2025 | Taniguchi G. | Tsubouchi H. | 40825586 | rad51 h+, rad52 h+, rad54 h+ | |
| 2025 | Bakosova A. | Polakova S.B. | 40594858 | rad51 h+, rad52 h+ | |
| 2025 | Lu G. | Feng G. | 39789818 | rad51 h+, rad52 h+ | |
| 2024 | Qin B. | Feng G. | 38718864 | rad51 h- | FY18537 |
| 2023 | Mongia P. | Nakagawa T. | 37237082 | rad51 smt0 | |
| 2022 | Kishkevich A. | Whitby M.C. | 36435847 | rad51 h-, rad54 h-, most h+ or ur | MCW9013, MCW9700, MCW9723, MCW9725, |
| 2022 | Disbennett W.M. | Petreaca R.C. | 35622511 | rad51 h-, rad52 h- | RCP19, RCP176 |
| 2022 | Vines A.J. | King M.C. | 35080989 | rad51 h- | MKSP2709 |
| 2021 | Su J. | Nakagawa T. | 34292936 | rad51 and rad52 in h+ or h-smt0 | |
| 2021 | Misova I. | Polakova S.B. | 33511417 | rad51 h- | S7068 |
| 2021 | Yamamoto I. | Ishikawa F. | 34520548 | rad51 h-? | IY4890, IY4899, TM2782 |
| 2021 | Watson A.T. | Carr A.M. | 34201031 | rad52 h+ | |
| 2020 | Kramarz K. | Lambert S.A.E | 33159083 | rad51 h-, rad52 h- | KK58, KK59, KK172 |
| 2020 | Argunhan B. | Iwasaki H. | 32204793 | rad51 smt0 | |
| 2020 | Matmati S. | Coulon S. | 32160539 | rad51 h+ | |
| 2020 | Dave A. | Humphrey T.C. | 31828313 | rad51 h- | TH2999, TH3034 |
| 2020 | Shetty M. | Noguchi E. | 31833215 | rad52 h+ | |
| 2019 | Yan Z. | Ira G. | 31542296 | rad52 h-, rad51 h- | YSK124, YSK180, YSK187, yZY031, and others |
| 2019 | Lucas B.E. | Petreaca R.C. | 31454903 | rad52 h-, rad51 h- | RCP71, RCP81, RCP258, RCP288, RCP377 |
| 2019 | Wong I.N. | Whitby M.C. | 31855181 | rad51 h-, rad51 mat undetected | MCW9496, (MCW8296?) |
| 2019 | Hardy J. | Lambert S. | 31584934 | rad52 h+, rad52 h- smt-0 | |
| 2019 | Tamang | Whitby M.C. | 31149897 | rad51 h+ | |
| 2017 | Teixeira-Silva A | Lambert S.A.E | 29215009 | rad51 h+ /smt-0, rad52 h+/smt-0 | |
| 2017 | Zafar F. | Nakagawa T. | 28977643 | rad51 h+, rad52 h+, rad54 h+ | |
| 2017 | Noguchi C. | Noguchi E. | 27687866 | rad52 h- | Y4753, Y5110 |
| 2017 | Morrow C.A. | Whitby M.C. | 28586299 | rad51 h+, rad52 h+ | |
| 2017 | Ait Saada A. | Lambert S.A.E | 28475874 | rad51 h+ /smt-0, rad52 h+/smt-0 | |
| 2016 | Onaka A.T. | Nakagawa T. | 27697832 | rad51, rad52, rad54 h+ or h- smt-0 | |
| 2016 | Zabrady K. | Palecek J.J. | 26446992 | rad51 h- | FY18537 |
| 2016 | Polakova S. | Gregan J. | 27304859 | rad51 h- | JG17507 |
| 2016 | Aronica | Humphrey T.C. | 26682798 | rad51 h- | TH2801 |
| 2016 | Ohno Y | Ishii K. | 26433224 | rad51 h- smt0, rad52 h- smt0 | |
| 2016 | Gadaleta C. | Noguchi E. | 26990647 | rad52 h- | Y3784 |
| 2016 | Callegari A.J. | Kelly T.J. | 26652183 | rad51 h- smt0 | |
| 2015 | Nguyen M.O. | Whitby M.C. | 25806683 | rad51 h+ /h- smt-0, rad52 h+/h- smt-0 | |
| 2015 | Audry J. | Coulon S. | 26041456 | rad51 h+ | |
| 2014 | Pietrobon V. | Lambert S.A.E | 25313826 | rad51 h+ /h- smt-0, rad52 h+/h- s | |
| 2014 | Tsutsui Y | Iwasaki H. | 25165823 | rad51 h+ | |
| 2014 | Swartz | King M.C. | 24943839 | rad51 h- | FY18537,... |
| 2014 | Nakano A. | Ueno M. | 24469396 | rad51 h- | FY18537,.... |
| 2014 | Stanescu R.S. | Petrescu-Danila E. | 24741789 | rad51 h- | AMC60 |

**Table S2** - *S. pombe* strains used in this study

| strain # | genotype | Used in | source |
|---|---|---|---|
| AMC501 | *h- ade6-704 leu1-32 ura4-D18* | Fig. 1C; 3B; 4A,C; 5A-B; 6A; S2 | A.M. Carr |
| yPK672 | *h+ rad51::KanMX4 ade6-216 leu1-32 ura4-D18* | Fig. 1C; 2A-C; 6B | Bioneer library |
| yPK614 | *h+ rad52::KanMX4 ade6-216 leu1-32 ura4-D18* | Fig. 1C; 2A-C | Bioneer library |
| SP25 | *h+ rad54::KanMX ade6-704 leu1-32 ura4-D18* | Fig. 1C; 2A-C | S. B.-Polakova |
| yPK546-8C | *h- smt-0 ade6-704 leu1-32 ura4-D18* | Fig. 2A | This study |
| yPK530-4C | *h- swi1::leu+ ade6-704 leu1-32 ura4-D18* | Fig. 2B | This study |
| yPK459-3D | *h- fml1::NatMX4 ade6-704 leu1-32 ura4-D18* | Fig. 2C | This study |
| yPK627-1A | $h^{90}$ *fml1::NatMX4 ade6-704 leu1-32 ura4-D18* | Fig. 2D | This study |
| AMC503 | *h+ ade6-704 leu1-32 ura4-D18* | File S1 | A.M. Carr |
| FY18537 | *h- rad51::hygr leu1-32 ura4-D18* | File S1 | NBRP |
| FY18725 | *h+ (rad51::his3) ade7-152 his3-D1* | text | NBRP |
| FY20076 | *h(-)+ rad51::ura4+* | text | NBRP |
| yPK444 | $h^{90}$ *ade6-704 leu1-32 ura4-D18* | File S1; Fig. 3B; 4A,C; 5A-B; 6A; S2 | S. B.-Polakova |
| yPK795-8B | *h+ ade6-216 leu1-32 ura4-D18* | Fig. 3B; 4A,C; 5A-B; 6A; S2 | This study |
| yPK804-1A | *h+ clr4::KanMX6 ade6-704 leu1-32 ura4-D18* | Fig. 4A,C; 5A-B; 6A-B; S2 | This study |
| yPK799-10 | *h+ mat1-cenHΔ::ura4+ ade6-704 leu1-32 ura4-D18* | Fig. 4C; 5B; 6A; S2 | This study |
| yPK402 | *h+ atf1::KanMX4 ade6-216 leu1-32 ura4-D18* | Fig. 4C; 5B; 6A; S2 | Bioneer library |
| yPK803-1A | *h+ mat1-cenHΔ::ura4+ atf1::KanMX4 ade6-704 leu1-32* | Fig. 4C; 5B; | This study |
| FY11920 | *h+ clr6-1 leu1-32 his2* | Fig. 6A; S2 | NBRP |
| SP435 | *h+ dcr1::KanMX4 leu1-32 his2* | Fig. 6A; S2 | S. B.-Polakova |
| JB718 | *h- clr4::kanMX6 ade6-M210 leu1-32 ura4-D18, his3-D1* | Fig. 6B | J. Bahler |
| yPK765-5C | *h- smt-0 rad51::KanMX4 ade6-216 leu1-32 ura4-D18* | Fig. 6B | This study |

**Table S3**. Primers used in this study

| primer | | sequence 5' to 3' | Used in |
|---|---|---|---|
| oPK986 | Mat1-F | GAAGGAAAAATATTGGAAGAGGTAG | left-M - Fig. 3B; S2 |
| oPK987 | Mat-M-start-R | AAATTCTAACATCAAATTACAACTTAAACG | |
| oPK986 | Mat1-F | GAAGGAAAAATATTGGAAGAGGTAG | left-P - Fig. 3B; 4A; S2 |
| oPK981 | Mat-P-R | GGACGTTAGGAGACAAATTGCC | |
| oPK984 | Mat-M-F | CATGGATTTTACTGCCCTGATTC | right-M - Fig. 3B; 6A |
| oPK979 | Mat1-R | AAGGTAGAAGGGCGCACACA | |
| oPK985 | Mat-P-F | GGCCAATTCTACGAAGTTTTAAGG | right-P - Fig. 3B; 4A; 6A |
| oPK979 | Mat1-R | AAGGTAGAAGGGCGCACACA | |
| oPK1123 | rga7-F | CTCTGCGAGTCAAACACCCT | *rga7* - Fig. 4A; 5A |
| oPK1124 | rga7-R | ATGCTTGATGGAGGCGTGAA | |
| oPK1045 | NCRNA388-F | GACCCTTCGCTATCATCCCG | *NCRNA.388* - Fig. 4A,C; 5A,B |
| oPK1046 | NCRNA388-R | AAAGTTGTGTGGAGAGGCGA | |
| oPK1047 | SPBC23G7.10c-F | ATTCAATTGGCGCATGCTGG | *SPBC23G7.10c* - Fig. 4A,C; 5A,B |
| oPK1048 | SPBC23G7.10c-R | GTTGAGCGTGGTTTTCGTCC | |
| oPK1059 | mag2-qPCR-F | CAGACGCTGCCACAAATTCA | *mag2* - Fig. 4A; 5A |
| oPK1060 | mag2-qPCR-R | GCAGCCTCTGCAACAATGTG | |
| oPK1117 | SPBC23G7.13c-F | GACGACTGGTGCAAACAACC | *SPBC23G7.13c* - Fig. 4A |
| oPK1118 | SPBC23G7.13c-R | TTGCAAACGTGGCCAACAAA | |
| oPK984 | Mat-M-F | CATGGATTTTACTGCCCTGATTC | *mat3-M* - Fig. 4A |
| oPK1143 | Mat3-M-R | CCCAAAAGACAGACTATCGGC | |
| oPK966 | Mat1-Mc-F | CAGCGGGTTCCCCTATTTCC | *mat1-Mc* - Fig. 5A |
| oPK967 | Mat1-Mc-R | TGAGGTCTTGGCAGTTGTGC | |
| oPK971 | Mat1-Pc-F | AGTATGCGCTCTAACTTGGCA | *mat1-Pc* - Fig. 5A |
| oPK970 | Mat1-Pc-R | GGTGCTTCAGCCAAATGCTC | |
| oPK1115 | rpt6-F | CCTGCTCTTCTTCGTCCAGG | *rpt6* - Fig. 5A |
| oPK1116 | rpt6-R | TTTAGCTCGGCACCACTAGC | |
